# Breaking barriers: The effect of protected characteristics and their intersectionality on career transition in academics

**DOI:** 10.1101/593905

**Authors:** Klara M. Wanelik, Joanne S. Griffin, Megan Head, Fiona C. Ingleby, Zenobia Lewis

## Abstract

In the past decade the scientific community has been trying to tackle the historical underrepresentation of women in science and the fact that gender can constitute a barrier to career success. However, other characteristics, such as being of an ethnic minority or coming from an under-privileged background, have received less attention. In this study we find that ethnicity and socioeconomic status impact detrimentally on career progression in early career scientists, despite the fact that gender is more likely to be reported as a barrier. Our data suggest we need to widen the discussion regarding diversity and equality in science to incorporate potential barriers to career success in addition to gender.

**Abstract:** The academic disciplines of Science Technology Engineering and Mathematics (STEM) have long suffered from a lack of diversity. While in recent years there has been some progress in addressing the underrepresentation of women in STEM subjects, other protected characteristics have received less attention. In this study, we survey early career scientists in the fields of ecology, evolutionary biology, behaviour, and related disciplines. We (i) quantitatively examine the effect of protected characteristics and their intersectionality on career transition, and (ii) provide practical suggestions, based on the qualitative responses of those surveyed, for overcoming some of the barriers we identified. We found that socioeconomic background and ethnicity impacted negatively on the quantitative measures of career progression we examined. Respondents that were female, LGBT, and from a lower socioeconomic background were more likely to report having faced a barrier, and the most frequent barrier named was related to gender. Our results suggest that respondents may have felt more confident discussing the experiences they have had related to their gender, potentially because there is now widespread discourse on this subject. However, respondents were less likely to discuss barriers they have faced in relation to ethnicity and socioeconomic status, despite the fact that the data indicates these are more detrimental to career progression. This may reflect the fact that these characteristics have received less attention, and are therefore deemed more sensitive. We hope that this study will stimulate wider discussion, and help to inform strategies to address the underrepresentation of minority groups in STEM subjects.

## Introduction

It is now widely recognised that diversity in the workforce is beneficial. In the private sector, numerous studies have shown that companies that are more diverse in terms of gender and/or ethnicity, exhibit greater performance in terms of outputs, growth, and financial gains (e.g. 1, 2, 3, 4). To date, the impacts of diversity in academia have been less well studied, which represents a significant gap in the literature (5). However, studies suggest that higher departmental diversity is related to higher placing in institutional rankings (6), gender-diverse collaborative groups produce higher quality science, as reviewed by peers (7), and ethnically diverse groups produce papers with higher scientific impact (8).

The disciplines of Science Technology Engineering and Mathematics (STEM) have historically suffered from a lack of diversity. In the UK for example, according to the 2015-16 Higher Education Statistics Agency (HESA) data, academics working in STEM subjects were 41.4% female, while 51% of the national population are women. Where the data are broken down according to role, women are represented much less than men in senior positions, even though at undergraduate level female students outnumber male students; this loss of female representation with academic progression has been dubbed the ‘leaky pipeline’ effect (9, 10). Internationally the figures are even more stark, with only 32% of researchers in STEM being female in Western Europe and North America, and 29% worldwide (11). In recent years the academic community has made some progress in addressing the historic underrepresentation of women in STEM subjects. Initiatives such as the UK-based Athena SWAN Charter (https://www.ecu.ac.uk/equality-charters/athena-swan/) and USA-based ADVANCE Programme run by the National Science Foundation (https://www.nsf.gov/crssprgm/advance/) have gained momentum. Professional Societies are investigating ways to increase visibility of women at conferences (e.g. 12), and journals are considering the equity of their publication processes (e.g. 13). Targeted training programmes such as the Aurora Leadership Programme for Women in Science (https://www.lfhe.ac.uk/en/programmes-events/equality-and-diversity/aurora/) are supporting women in their academic progression. However, despite these measures, studies suggest there is still progress to be made to promote gender equality in STEM; for example a recent analysis suggested it will take generations to achieve gender parity (14).

Minorities identifying with other characteristics, also protected under the Equality Act 2010, including race, disability, sexual orientation, and age, have received even less attention. There has been very little discussion with regards to the effect of coming from a protected group, despite evidence that this can have a strong effect on the probability of retaining an academic career (e.g. 15, 16, 17). Indeed, according to 2015-16 UK HESA statistics, STEM academics were 10.3% non-white, and 0.03% disabled, in contrast with the student demographics for the same time period, of 21%, and 11% respectively; in 2016-17, only 0.6% of UK professors were black. Individuals from protected groups can face a multitude of barriers, including financial worries, and negative perceptions of their own academic career success (18). Likelihood of promotion is lower than for non-minority groups (e.g. 19), and they may be less able to access voluntary positions and internships to gain experience and training (reviewed in 20).

Where individuals identify with more than one protected characteristic, the challenges faced by individuals can be further compounded, a situation referred to as the ‘double bind’ (21, 22). For example, ethnic minority female academics are more likely to suffer from self-doubt, and are more likely to experience challenges to their authority, compared to white male and female academics (reviewed in 23), while female LGBT students have been shown to exhibit the lowest feeling of belonging to their academic community compared to other groups (24). These examples demonstrate the importance of an intersectional approach, considering protected characteristics together, rather than in isolation.

In the present study, we present data collected from a survey with 205 respondents, all early career researchers (within ten years of completing their PhD) in the fields of ecology, evolutionary biology, behaviour, and related disciplines (the fields of the authors). We use these data to (i) examine the effect of six characteristics (ethnicity, age, sexual orientation, sex, socioeconomic background and disability) and their intersectionality on career transition of academics in these fields, and (ii) provide practical suggestions, based on the experience of respondents, for overcoming some of the barriers identified in (i). Although socioeconomic background is not a characteristic protected by the Equality Act 2010, we included this in our study as it has been suggested that financial barriers can make it harder for early career researchers to progress in science (25). We hope that this study will help to inform strategies to address the underrepresentation of minority groups in STEM subjects.

## Methods

To obtain information on the barriers faced by early career scientists in the fields of ecology, evolutionary biology, behaviour, and related disciplines, we conducted an online survey (Table S1), hosted by SurveyMonkey, Inc. (USA). The link to the survey was communicated via social media and email, via Evolutionary Directory (EvolDir), with a simple title ‘STEM survey’. We left the survey open for 3 weeks during which we received 205 anonymous responses. Ethical approval to collect the responses was required and granted by the University of Liverpool (application reference number 2229). The responses were collected anonymously and voluntarily. We specified that respondents should be early career scientists within a maximum of ten years of completing their PhD. For transparency, a set of questions was outlined in a research plan prior to analysis. This plan also included detailed methods for answering each of these questions, which are described below (original research plan available at: https://github.com/kwanelik/Breaking-barriers).

### Summary of respondent demographics

Respondents to our survey were geographically diverse, with good numbers having completed their PhD in the US & Canada (*n* = 64), Europe (*n* = 48) or Australia & New Zealand (*n* = 20). Other regions included Scandinavia (n = 7) and Central/South America (n = 5). The modal age of respondents was 30-34 and the modal age upon PhD submission was 25-29. The majority of respondents were on research-only contracts (*n* = 76; as opposed to teaching-only or research and teaching combined contracts) and were not on a permanent contract at the time of completing the survey (*n* = 107). Approximately equal numbers of respondents reported either having faced barriers (*n* = 71) or not (*n* = 60). A full breakdown of numbers of respondents in relation to the protected characteristics of interest are provided in Table 1.

**Table 1.** A breakdown of respondents in relation to the protected characteristics of interest: ethnicity, age, sexual orientation, sex, disability and socioeconomic background

| Protected characteristic | N |
| --- | --- |
| **Ethnic group** |  |
| Black, Asian and minority ethnic (BAME) | 14 |
| Latino Hispanic | 15 |
| White | 154 |
| Other | 5 |
| **Age current/PhD** |  |
| 18-24 | 12 |
| 25-29 | 35 |
| 30-34 | 77 |
| 35-39 | 38 |
| 40-44 | 21 |
| 45-49 | 4 |
| 50+ | 1 |
| **Sexual Orientation** |  |
| Straight | 155 |
| LGBT | 24 |
| Prefer not to answer | 9 |
| **Sex** |  |
| Females | 123 |
| Male | 63 |
| Other | 2 |
| **Disability** |  |
| Yes | 12 |
| No | 175 |
| Prefer not to answer | 1 |
| **Socioeconomic background** |  |
| Lower Yes | 45 |
| Lower No | 139 |
| Prefer not to answer | 3 |

### Quantitative analysis

#### Data cleaning

In some cases, answers given by respondents were ambiguous. Where respondents included a lower limit (e.g. number of postdoc applications made before being awarded a position = “100+”), this was used. Where they included a range (e.g. “15-20”) a mean value was used. Where they included an approximate figure (e.g. “approx. 100”) these were treated the same as exact figures. Sixteen responses were discounted due to ambiguous answers which could not be confidently interpreted. The minimum timescale was taken to be one, so any timescales less than one year were rounded up. Outliers were defined as those data points lying more than three standard deviations from the mean, and were removed prior to analysis. Some groups with very low representation e.g. other/prefer not to answer also had to be removed due to problems with model convergence. Individuals with missing values for any independent variables of interest were also excluded. Final sample size for each analysis are included in Table 2.

**Table 2.** Table detailing the global models for each of the questions

| **Response variable** | ***n*** | **GLM Error family** | **Link function** | **Global model** |
| --- | --- | --- | --- | --- |
| ***Publication records*** |  |  |  |  |
| No. first author papers | 144 | Negative Binomial | Log | Sex, Sexual orientation, Ethnic group, Socioeconomic background, Disability, Age PhD, Year PhD, Sex x Sexual orientation, Sex x Ethnic group, Sex x Disability |
| No. other author papers | 144 | Negative binomial | Log | Sex, Sexual orientation, Ethnic group, Socioeconomic background, Disability, Age PhD, Year PhD, Sex x Sexual orientation, Sex x Disability |
| ***Job application success*** |  |  |  |  |
| No. postdoc application | 126 | Negative binomial | Log | Sex, Sexual orientation, Ethnic group, Socioeconomic background, Disability, Age PhD, Year PhD, Total publications PhD, Sex x Sexual orientation, Sex x Ethnic group, Sex x Age PhD, Sex x Total publications PhD |
| ***Types of contract*** |  |  |  |  |
| Research vs. Teaching & research | 111 | Binomial | Logit | Sex, Sexual orientation, Ethnic group, Socioeconomic background, Disability, Year PhD, Total publications PhD, Total postdocs, Permanent or not, Sex x Sexual orientation, Sex x Ethnic group, Sex x Disability, Sex x Total publications PhD |
| Permanent contract or not | 139 | Binomial | Cloglog | Sex, Age current, Sexual orientation, Ethnic group, Socioeconomic background, Disability, Total postdocs, Total publications PhD, Year PhD, Sex x Sexual orientation, Sex x Ethnic group, Sex x Disability, Sex x Total postdocs, Sex x Total publications PhD |
| ***Grant success*** |  |  |  |  |
| No. grant applications | 122 | Negative binomial | Log | Sex, Sexual orientation, Ethnic group, Socioeconomic background, Disability, Year PhD, Total publications PhD, Total postdocs, Sex x Sexual orientation, Sex x Ethnic group, Sex x Disability, Sex x Total publications PhD |
| ***Reported barriers*** |  |  |  |  |
| Reported barrier or not | 133 | Binomial | Logit | Sex, Sexual orientation, Ethnic group, Socioeconomic background, Disability, Year PhD |

#### Statistical modelling

Generalised linear models (GLMs) were used to look for associations between an individual’s protected characteristics and (i) publication record, (ii) job application success (iii) type of contract (iv) grant success, and (v) reported barriers. Model specifications (error distributions and link functions) are detailed in Table 2. All analyses were conducted in R version 3.4.4 (26). All models included the six protected characteristics as fixed effects (ethnicity, age, sexual orientation, sex, socioeconomic background and disability). Due to limited sample sizes, only interactions between sex and the other protected characteristics were included (where possible; see Table 2). We collected data on the country of PhD, with an aim to account for any geographical variation in our analyses. However, we did not include this variable in our final analyses, primarily to avoid overfitting (models including country as a random effect did not converge). This variable was also deemed to be of limited use, as a result of the geographic mobility of most academics. Year of PhD completion was included as a fixed effect in all models to account for any temporal autocorrelation. We originally planned for this variable to be included as a random effect but mixed effect models did not converge (likely because of the limited sample size). Therefore, years were combined into five bins of approximately equal size (2007-2009, 2010-2011, 2012-2013, 2014-15, 2016-2017) and included as a fixed effect. Other additional fixed effects included total publications at the time of PhD submission, total number of postdocs completed, whether or not on a permanent contract and interactions between these variables and sex (see Table 2). All fixed effects within a model were checked for collinearity by computing generalised variance inflation factors (GVIFs). Any fixed effects excluded on these grounds are detailed below. All models were checked for normal and homoscedastic residuals.

Sets of candidate models were generated from each global model, which included all of the fixed and interaction terms of interest (Table 2) using the MuMIn package (27). All candidate models were then ranked on relative fit using the Akaike information criteria corrected for small sample size, AICc (28). Those with a ΔAICc < 2 relative to the lowest value were considered to be equally supported as the best models to explain the data (top models). Effect sizes, unconditional standard errors and estimated *p*-values were obtained by averaging across this set of top models using the zero method (29). Model averaging was carried out due to the lack of a single best model. All reported effect sizes are on a transformed scale. Where two or more numeric variables were present in an averaged model, these were standardized using two SD (30) to make them directly comparable. The relative importance of a variable was taken to be the sum of the Akaike weights of the top models in which it was found (27). Variables that appear in one or more top model, but are not significant, are still reported. Even though there is no evidence for such variables affecting the response, they are still considered useful in predicting point estimates (31).

##### Publication record

We looked for associations between protected characteristics and the number of first- and other-author papers published upon PhD submission. The number of first- and other-author papers upon PhD submission were modelled separately. The number of other-author papers was somewhat zero-inflated (with 34 respondents having no other-author publications upon PhD submission) causing overdispersion, but a GLM with a negative binomial error distribution achieved acceptable residuals.

##### Job application success

We looked for associations between protected characteristics and the number of applications made before commencing an advertised postdoc position or fellowship (combined). In a separate model, we also looked for associations between protected characteristics and the number of applications made for permanent positions, for those respondents with permanent contracts. We grouped BAME and Hispanic-Latino individuals together due to too few respondents with permanent contracts, and we discarded disability from the analysis as only one respondent with a permanent contract reported having a disability. For both of these models, because we were interested in relating success to effort, we excluded those respondents which gave the number of job applications, but later stated that they had not yet been successful in securing a job (whether postdoc or permanent position; *n* = 3).

##### Types of contract

We looked for associations between protected characteristics and the type of contracts respondents were on (either research only or teaching and research). As only 5 respondents were on teaching-only contracts these were excluded from the analysis. Age and year of PhD were found to be highly collinear with other variables (year of PhD GVIF = 3.3; age GVIF = 2.5) and were therefore both excluded from this analysis.

We also looked for associations between protected characteristics and whether or not an individual was on a permanent contract. The vast majority of respondents (73%) were not on a permanent contract causing the data to be highly zero-inflated, but this did not violate the assumptions of the binomial GLM. Age was binned into two main groups: < 34 and > 34 years due to problems with model convergence. Year of PhD was found to be collinear with other variables (GVIF = 2.9) and was therefore excluded from this analysis.

##### Grant success

We looked for associations between protected characteristics and the number of grant applications made. We excluded small grants (e.g. travel grants), by specifically asking respondents about grants which included their salary. Age was found to be collinear with other variables (GVIF = 4.3) and was therefore excluded from this analysis.

##### Reported barriers

We looked for associations between protected characteristics and whether or not an individual reported facing barriers to their identity. Due to problems with model convergence, no interaction terms were included. Age was again found to be highly collinear with other variables (GVIF = 12.3) and so was excluded.

### Qualitative analysis

All free text answers from respondents on (i) the most important barriers they have faced, and (ii) how they overcame these barriers were analysed using the text mining (tm) package (32, 33) in R version 3.4.4 (26). Briefly, text was transformed and cleaned (removing all numbers, punctuation and stopwords). Then, text-stemming was performed and frequencies of root words were calculated. The most frequently used words are reported below.

## Results

### (i) Identification of barriers

#### Publication record

No protected characteristics were found to have a significant effect on the number of first-author papers published upon submission of PhD. However, disability, socioeconomic background and ethnicity were all present in the top models, suggesting that they may be useful predictors. Socioeconomic background appeared most frequently (three out of five top models; Table S2), but was not found to be significant (*p* = 0.48; Table 3).

**Table 3.**
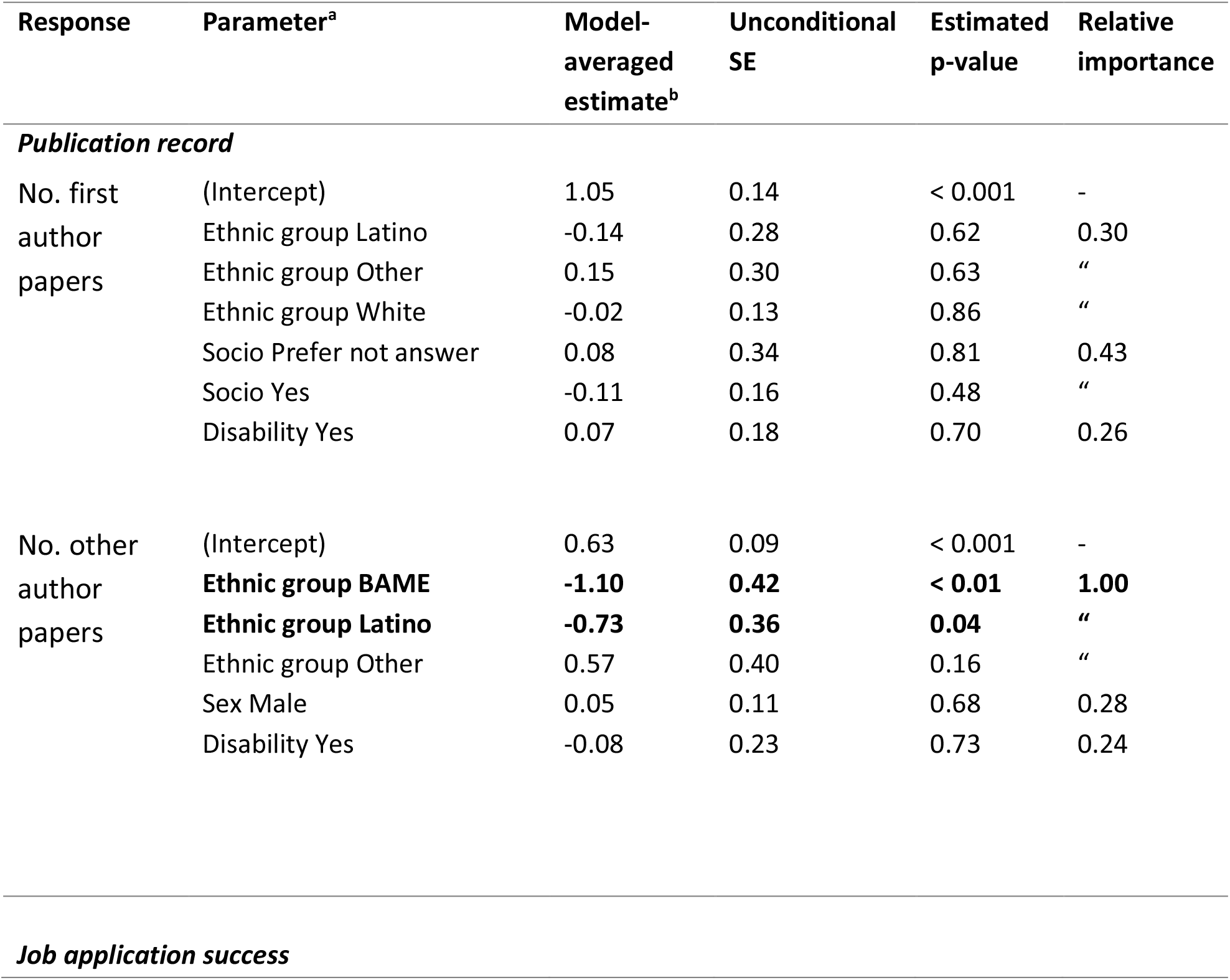
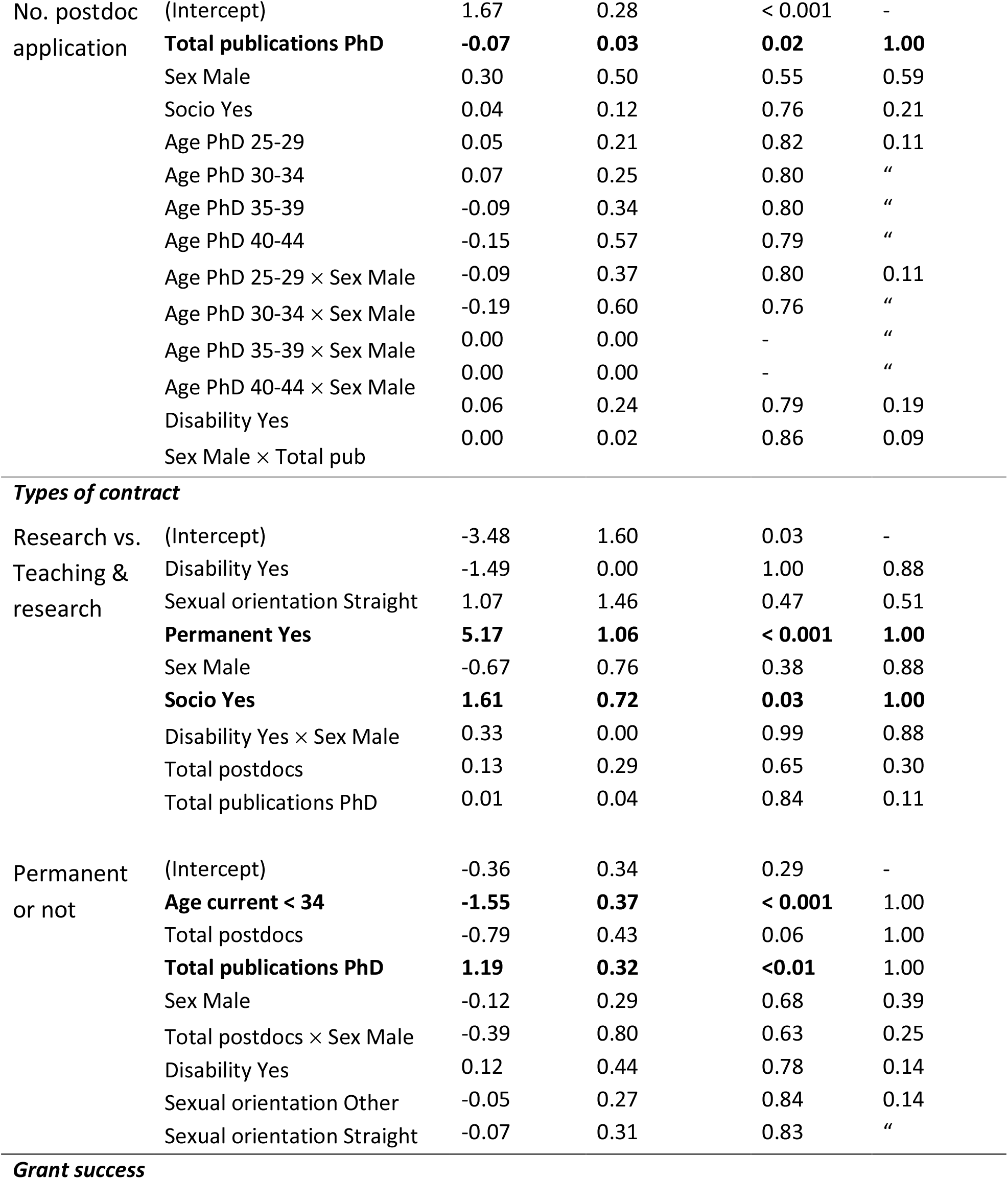
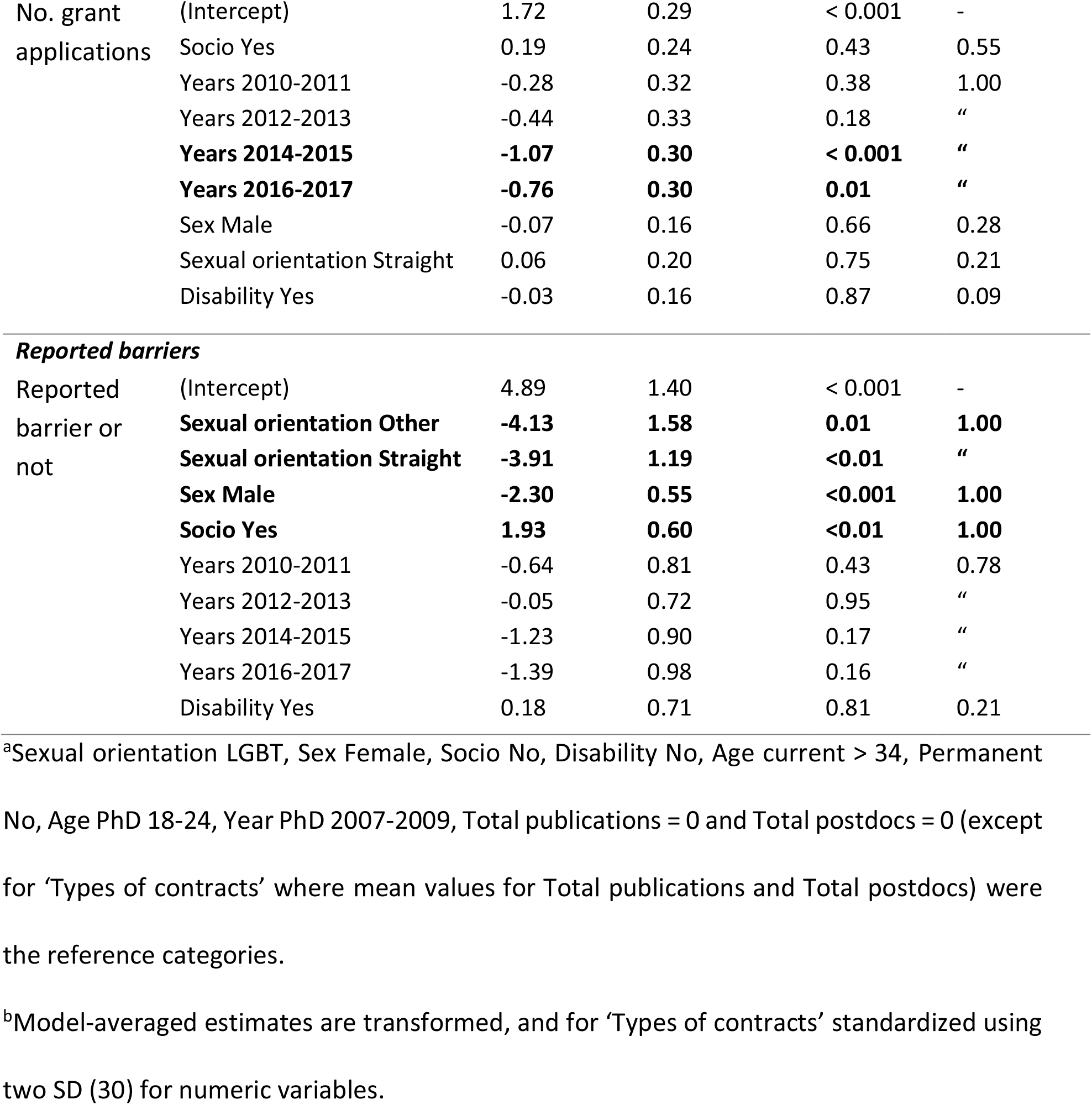
Model-averaged, transformed parameter estimates, unconditional standard errors, estimated *p*-values and relative importance of predictors of (i) Number of first author papers on PhD submission, (ii) Number of other author papers on PhD submission, (iii) Number of postdoc applications, (iv) Research vs. Teaching & research contract, (v) Permanent contract or not, (vi) Number of grant applications, and (vii) Reported barrier or not. Significant effects shown in bold.

| Response | Parameter<sup>a</sup> | Model-averaged<br>estimate<sup>b</sup> | Unconditional<br>SE | Estimated<br>p-value | Relative<br>importance |
| --- | --- | --- | --- | --- | --- |
| ***Publication record*** |  |  |  |  |  |
| No. first<br>author<br>papers | (Intercept) | 1.05 | 0.14 | < 0.001 | - |
|  | Ethnic group Latino | -0.14 | 0.28 | 0.62 | 0.30 |
|  | Ethnic group Other | 0.15 | 0.30 | 0.63 | “ |
|  | Ethnic group White | -0.02 | 0.13 | 0.86 | “ |
|  | Socio Prefer not answer | 0.08 | 0.34 | 0.81 | 0.43 |
|  | Socio Yes | -0.11 | 0.16 | 0.48 | “ |
|  | Disability Yes | 0.07 | 0.18 | 0.70 | 0.26 |
| No. other<br>author<br>papers | (Intercept) | 0.63 | 0.09 | < 0.001 | - |
|  | **Ethnic group BAME** | **-1.10** | **0.42** | **< 0.01** | **1.00** |
|  | **Ethnic group Latino** | **-0.73** | **0.36** | **0.04** | “ |
|  | Ethnic group Other | 0.57 | 0.40 | 0.16 | “ |
|  | Sex Male | 0.05 | 0.11 | 0.68 | 0.28 |
|  | Disability Yes | -0.08 | 0.23 | 0.73 | 0.24 |
| ***Job application success*** |  |  |  |  |  |
| No. postdoc application | (Intercept) | 1.67 | 0.28 | < 0.001 | - |
|  | **Total publications PhD** | **-0.07** | **0.03** | **0.02** | **1.00** |
|  | Sex Male | 0.30 | 0.50 | 0.55 | 0.59 |
|  | Socio Yes | 0.04 | 0.12 | 0.76 | 0.21 |
|  | Age PhD 25-29 | 0.05 | 0.21 | 0.82 | 0.11 |
|  | Age PhD 30-34 | 0.07 | 0.25 | 0.80 | " |
|  | Age PhD 35-39 | -0.09 | 0.34 | 0.80 | " |
|  | Age PhD 40-44 | -0.15 | 0.57 | 0.79 | " |
|  | Age PhD 25-29 × Sex Male | -0.09 | 0.37 | 0.80 | 0.11 |
|  | Age PhD 30-34 × Sex Male | -0.19 | 0.60 | 0.76 | " |
|  | Age PhD 35-39 × Sex Male | 0.00 | 0.00 | - | " |
|  | Age PhD 40-44 × Sex Male | 0.00 | 0.00 | - | " |
|  | Disability Yes | 0.06 | 0.24 | 0.79 | 0.19 |
|  | Sex Male × Total pub | 0.00 | 0.02 | 0.86 | 0.09 |
| ***Types of contract*** |  |  |  |  |  |
| Research vs. Teaching & research | (Intercept) | -3.48 | 1.60 | 0.03 | - |
|  | Disability Yes | -1.49 | 0.00 | 1.00 | 0.88 |
|  | Sexual orientation Straight | 1.07 | 1.46 | 0.47 | 0.51 |
|  | **Permanent Yes** | **5.17** | **1.06** | **< 0.001** | **1.00** |
|  | Sex Male | -0.67 | 0.76 | 0.38 | 0.88 |
|  | **Socio Yes** | **1.61** | **0.72** | **0.03** | **1.00** |
|  | Disability Yes × Sex Male | 0.33 | 0.00 | 0.99 | 0.88 |
|  | Total postdocs | 0.13 | 0.29 | 0.65 | 0.30 |
|  | Total publications PhD | 0.01 | 0.04 | 0.84 | 0.11 |
| Permanent or not | (Intercept) | -0.36 | 0.34 | 0.29 | - |
|  | **Age current < 34** | **-1.55** | **0.37** | **< 0.001** | **1.00** |
|  | Total postdocs | -0.79 | 0.43 | 0.06 | 1.00 |
|  | **Total publications PhD** | **1.19** | **0.32** | **<0.01** | **1.00** |
|  | Sex Male | -0.12 | 0.29 | 0.68 | 0.39 |
|  | Total postdocs × Sex Male | -0.39 | 0.80 | 0.63 | 0.25 |
|  | Disability Yes | 0.12 | 0.44 | 0.78 | 0.14 |
|  | Sexual orientation Other | -0.05 | 0.27 | 0.84 | 0.14 |
|  | Sexual orientation Straight | -0.07 | 0.31 | 0.83 | " |
| ***Grant success*** |  |  |  |  |  |
| No. grant applications | (Intercept) | 1.72 | 0.29 | < 0.001 | - |
|  | Socio Yes | 0.19 | 0.24 | 0.43 | 0.55 |
|  | Years 2010-2011 | -0.28 | 0.32 | 0.38 | 1.00 |
|  | Years 2012-2013 | -0.44 | 0.33 | 0.18 | " |
|  | **Years 2014-2015** | **-1.07** | **0.30** | **< 0.001** | **"** |
|  | **Years 2016-2017** | **-0.76** | **0.30** | **0.01** | **"** |
|  | Sex Male | -0.07 | 0.16 | 0.66 | 0.28 |
|  | Sexual orientation Straight | 0.06 | 0.20 | 0.75 | 0.21 |
|  | Disability Yes | -0.03 | 0.16 | 0.87 | 0.09 |
| **Reported barriers** |  |  |  |  |  |
| Reported barrier or not | (Intercept) | 4.89 | 1.40 | < 0.001 | - |
|  | **Sexual orientation Other** | **-4.13** | **1.58** | **0.01** | **1.00** |
|  | **Sexual orientation Straight** | **-3.91** | **1.19** | **<0.01** | **"** |
|  | **Sex Male** | **-2.30** | **0.55** | **<0.001** | **1.00** |
|  | **Socio Yes** | **1.93** | **0.60** | **<0.01** | **1.00** |
|  | Years 2010-2011 | -0.64 | 0.81 | 0.43 | 0.78 |
|  | Years 2012-2013 | -0.05 | 0.72 | 0.95 | " |
|  | Years 2014-2015 | -1.23 | 0.90 | 0.17 | " |
|  | Years 2016-2017 | -1.39 | 0.98 | 0.16 | " |
|  | Disability Yes | 0.18 | 0.71 | 0.81 | 0.21 |

<sup>a</sup>Sexual orientation LGBT, Sex Female, Socio No, Disability No, Age current > 34, Permanent No, Age PhD 18-24, Year PhD 2007-2009, Total publications = 0 and Total postdocs = 0 (except for 'Types of contracts' where mean values for Total publications and Total postdocs) were the reference categories.
<sup>b</sup>Model-averaged estimates are transformed, and for 'Types of contracts' standardized using two SD (30) for numeric variables.

The number of other-author papers published upon PhD submission differed significantly across ethnicity, with ethnicity appearing in all four top models. Both Black, African and minority ethnic (BAME; *p* < 0.01) and Hispanic-Latino individuals (*p* = 0.04) finished their PhD with approximately one less other-author publication than individuals of white ethnic background.

#### Job application success

We found a significant effect of the total number of papers on the number of postdoc applications (estimate = −0.07; *p* = 0.02), such that individuals with a greater combined number of first- and other-author papers make fewer postdoc applications, before obtaining a postdoc, than those with fewer publications. Age, sex, disability, socioeconomic background, sex × age and sex × total publications PhD all appeared in one or more top models (Table S2), but the number of applications did not differ significantly across these protected characteristics, nor their interactions (all *p* > 0.05; Table 3).

We have excluded question 3 of the original analysis plan from the results. This question asked whether the number of applications for permanent academic positions was associated with protected characteristics, for those individuals who had a permanent contract. Only 35 of the survey respondents had permanent contracts, therefore there was not enough data to reliably address this question. Of the analyses we were able to run, none of the candidate models in the set tested had ΔAICc < 2 relative to the top model, which included only the model intercept.

#### Types of contract

Socioeconomic background and job permanency were both present in all six top models for type of contract (research only or teaching and research; Table S2). As expected in the fields of ecology, evolutionary biology and behaviour, those who had permanent contracts were more likely to have teaching and research contracts than research-only contracts (estimate = 5.17; *p* < 0.001). Individuals from a lower socioeconomic background were also significantly more likely to have teaching and research contracts than research-only contracts, after accounting for job permanency (estimate = 1.61; *p* = 0.03). Other variables present in one or more top models were disability, sexual orientation, total publications PhD, total postdocs, sex and sex × disability but these had non-significant effects on the type of contract (all *p* > 0.05; Table 3).

The total number of publications had a positive effect on securing a permanent academic position (estimate = 1.19; *p* < 0.01). There was some evidence to suggest a significant negative association between age and securing a permanent position, such that individuals aged younger than 34 were less likely to secure a permanent position (estimate = −1.90; *p* < 0.001). Disability, sex, sexual orientation, total number of postdocs and sex × total number of postdocs were all present in one or more top models (Table S2; Table 3).

#### Grant success

There were no significant associations between the protected characteristics and the number of grant applications applied for (all *p* > 0.05). Socioeconomic background was present in four of seven top models but it was not significant (*p* = 0.43). As expected, there was a significant temporal effect, with those who submitted their PhD earlier having significantly more grant applications than those who handed in their PhDs later (e.g. Years 2007-2009 vs. Years 2014-2015; estimate = −1.07; *p* < 0.001; Table 3).

#### Reported barriers

Sex, sexual orientation and socioeconomic background were present in all three top models (Table S2; Table 3). LGBT individuals were significantly more likely to report facing a barrier than heterosexuals (estimate = 3.91; *p* < 0.01). Females were significantly more likely to report facing a barrier than males (estimate = 2.30; *p* < 0.001). Finally, individuals from a lower socioeconomic background were significantly more likely to report facing a barrier than those from a higher socioeconomic background (estimate = 1.93; *p* < 0.01).

Of the 71 free text responses received to the question ‘What do you feel was the most important [barrier]?’ the most frequently used words were related to sex; ‘woman/women’ (*n* = 17), ‘family’ (*n* = 10), ‘gender’ (*n* = 10), ‘female’, and ‘male’ (both, *n* = 7).

### (ii) Overcoming barriers

There were 55 free text responses to the question ‘If you were able to overcome the barrier stated in the previous question, how?’ Of these, approximately a third (*n* = 18) reported that they had not overcome their barriers and/or had left an institution, or academia all together, because of them. A number of words appeared at frequencies of three to five, which can be divided into main categories ‘people’ and ‘opportunities’. The ‘people’ category included phrases related to support, avoiding judgemental people, meeting people networking, associating with ‘high quality’ groups and senior allies, and mentoring. In terms of ‘opportunities’, suggestions that appeared several times were the importance of taking up opportunities, proving one’s self and skills, working hard, applying for many grants and positions, moving between institutions and asking for opportunities, both in negotiations, but also more generally. Other comments made with respect to opportunities, albeit less frequently, included the importance of perseverance, ensuring CVs are maintained well, participating in departmental activities, and seeking out paid work experience.

## Discussion

We used survey responses to address questions about the effect of protected characteristics on career transition, with a particular interest in widening our understanding of different types of protected characteristic and how these might interact. Although our results are complex, socioeconomic background and ethnicity had an impact on the measures of STEM career progression that we studied. We also found that multiple characteristics surveyed were somewhat important in predicting whether an individual reported facing a barrier, with sex, sexual orientation, and socioeconomic background all being particularly important.

Ethnicity was the main determinant of the number of publications obtained on finishing a PhD, although we cannot disentangle whether this effect is ethnicity per se, or country where PhD was awarded. Expectations of what is expected from a PhD differ between countries, and certain ethnicities are more likely to have undertaken their PhD in certain countries. There is some indication that socioeconomic background was also a predictor of numbers of papers. Although we found no direct relationship between the protected characteristics and job applications, we did find that having fewer publications on finishing a PhD translated into having to apply for more positions in order to secure a postdoctoral job. Similarly, having more publications translated into an increased likelihood of securing a permanent position. It’s therefore likely that the effect of protected characteristics on publication record could impact indirectly on future job applications and create a knock-on effect at a later career stage.

In addition to this, people from a lower socioeconomic background were more likely to be in teaching and research positions as opposed to research only positions. Possibly, teaching is viewed as a more ‘normal’ job, and therefore is more understandable and culturally acceptable to friends and family from non-academic backgrounds. Although we were unable to include teaching-only contracts in our analyses due to a small sample size, these positions can be associated with decreased job security and satisfaction, as many teaching positions at UK institutions tend to be fixed contract and do not have routes for promotion (34).

Socioeconomic status as an important determinant of career progression has been acknowledged elsewhere (e.g. 25), but the importance of financial support has received little widespread discussion in the wider STEM academic community, possibly because it is something of a sensitive topic. Given the precarious nature of STEM careers – often involving short-term contracts, and lengthy periods of unemployment and/or frequent geographic relocations between contracts – it is logical that familial wealth could prove a key determinant of whether an individual is able to progress to the next stage of their career. In addition, the culture of academia is one historically more associated with the upper classes, and individuals from a lower socioeconomic background are, perhaps, more likely to struggle with a lack of relatable role models, difficulty ‘fitting in’, and imposter syndrome (35, 36).

Financial barriers may be particularly relevant to ecology, evolution, behavioural ecology, and related disciplines, due to research in these fields often relying on field work. Experience with fieldwork can be key to career development; however, gaining this experience often requires undertaking voluntary internships, which may only be accessible to those from more privileged backgrounds (20).

Ethnicity has been reported elsewhere as having a negative impact on career progression (e.g. 37, 16), and our results are consistent with this. While overt discrimination based on ethnicity is no longer commonplace, ethnic minorities are more likely to experience institutional and cultural barriers to career progression (38, 39). Similarly to individuals from lower socioeconomic backgrounds, ethnic minority academics are less likely to have role models; more likely to suffer from imposter syndrome; more likely to lack a sense of ‘belonging’ in academia; and less likely to be promoted (reviewed in 40, 19, 41, 39).

We were interested in studying combinations of different protected characteristics to address the issue of career barriers being compounded for individuals that identify with more than one of the characteristics. As an example, ethnic minority women were the least represented group in UK academia in 2016-17, with only 25 black female professors out of 19,000 at that time (40), and reports on the experiences of this small group suggesting considerable barriers to career progression (39). Quantitative analysis of the experiences of such under-represented groups is often difficult due to sample size constraints, although we did find some evidence in our models for the interaction between sex and disability being a useful predictor of the type of contract.

In our qualitative analysis, we found that multiple protected characteristics studied were at least somewhat important in predicting if an individual a reported barrier. Respondents that were female, LGBT, or came from a lower socioeconomic background were the most likely to report having faced a barrier, and many of the responses cited gender as a barrier. Worryingly, almost half of the responses concerning ‘overcoming barriers’ were from respondents who stated they had left academia due to a barrier they had not been able to move past. Other respondents made suggestions for overcoming barriers which we divided into two main categories: ‘people’ and ‘opportunities’. With regards to ‘people’, several respondents mentioned mentoring, and indeed there is a wealth of literature that suggests that effective mentoring can be beneficial at all stages of a career (e.g. 43, 44, 4). Seeking senior allies and networking were also mentioned. Evidence suggests that the establishment of professional networks both inside and outside of the institution can be beneficial to career success (e.g. 45, 46), and diverse networks have been shown to be particularly advantageous (47). Conferences are an obvious route to networking, however, increasingly digital methods of building networks are available for women (e.g. SciSisters for female academics based in Scotland, http://www.chemicalimbalance.ed.ac.uk/scisister/ and 500 Women Scientists, https://500womenscientists.org/ which is worldwide), LGBT academics (e.g. The British Ecological Society LGBT+ Network https://www.britishecologicalsociety.org/membership-community/diversity/), and ethnic minority academics (e.g. the Twitter forum Minorities in STEM, https://twitter.com/minoritystem?lang=en).

Regarding the ‘opportunities’ category, many responses highlighted the importance of perseverance, working hard and always to a high standard, publishing as frequently as possible, applying to as many positions as possible, and ensuring CVs are maintained well. Proactive participation in departmental activities was deemed important by one respondent, while another suggested seeking out paid internships to gain work experience. These constructive suggestions are common themes in advice for overcoming bias in the workplace, although it is important to recognise the role of institutions in ensuring such opportunities are made accessible to all early-career STEM academics; institutional cultural change is needed to ensure that minority groups do not have to work harder to succeed or prove themselves.

It is worth highlighting that while sex featured strongly in our qualitative results, it was less significant in predicting career progression in our quantitative data. Our data do not allow us to determine whether the respondents had been successful in overcoming the gender barrier in terms of their career progression, nor whether this is representative of the wider community. It is possible that respondents felt more confident discussing gender in the free text comments as there is now a widespread narrative with regards to women in science. In contrast, the other characteristics, such as socioeconomic status and ethnicity, have received less attention, and so potentially people view these as more sensitive, and are less comfortable expressing their opinions.

In summary, our quantitative analyses suggested that socioeconomic status and ethnicity were important barriers to STEM career progression, with sexual orientation and gender also appearing important, and our qualitative analyses showed that gender was reported most frequently as a barrier. We find it worrying that gender is still deemed a significant obstacle to career success, suggesting that UK initiatives such as Athena SWAN have much work still to do. The importance of socioeconomic background is similarly worrying given increasing inequality and economic instability worldwide. Our models highlight a role of all protected characteristics in STEM careers, suggesting that ultimately there is a significant pool of the workforce who are struggling to access, retain, and succeed in a STEM academic career. As reported elsewhere, the challenges faced by individuals from protected groups are not only leading to a loss of diversity in the workplace, but also to the loss of talented individuals who could and should be meaningfully contributing to Higher Education (48).

Finally, we should be concerned that the picture may be even bleaker than it seems; our sampling method was unlikely to capture responses from many individuals who have already left academia as a result of the barriers they faced. In addition, the sample size for some groups, particularly those relating to disability and sexual orientation, were extremely small and this limited our ability to reliably analyse questions relating to these groups. Low representation of some protected groups in academia may be one reason why so much of the research carried out relies to some extent on qualitative rather than quantitative analysis.

Clearly, our community, and the STEM academic community more widely, has work to do. Community initiatives are making strides in breaking the barriers that face a substantial part of the population, but further support is needed at all levels. Nationally, we need to ensure that access to education and retention in the academic pipeline is inclusive to all. Individually, institutions have an important role in ensuring accessibility and inclusivity for student and staff hiring, retention and management. We hope this study stimulates open discussion and further research into this area.

## Author contribution statement

KMW, JSG, and ZL conceptualised the project. KMW, JSG, MH, FCI, and ZL designed the research. KMW, JSG, and ZL collected the data. KMW, JSG, and FCI analysed the data. All authors wrote the paper.

## Acknowledgements

This work was supported by a grant from the European Society of Evolutionary Biology Equal Opportunities Initiative Fund.

## Supplementary material

**Table S1:**
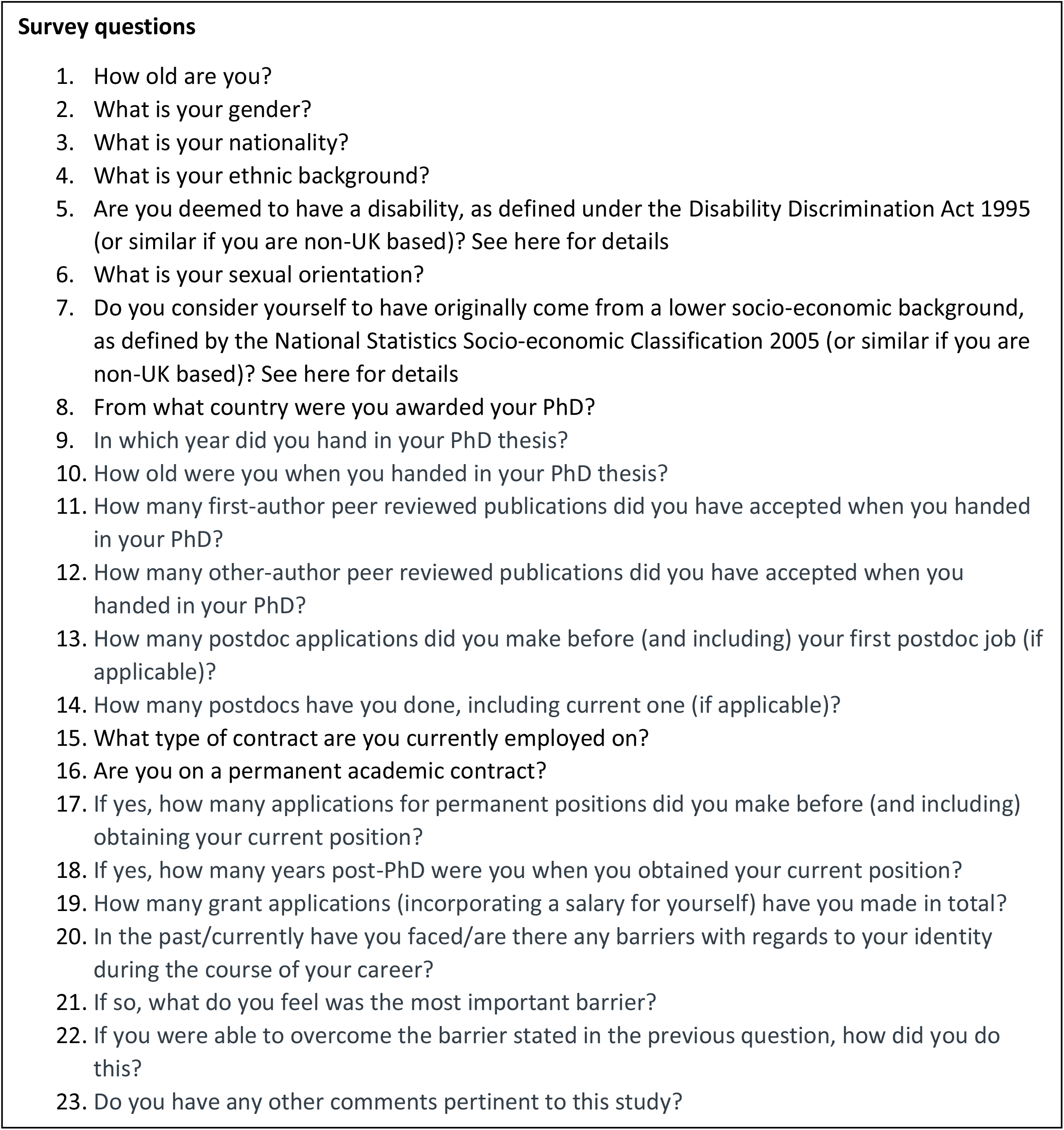
A table of survey questions

**Table S2.** Table detailing the top models for each of the questions

| **Response** | **Top models** |
| --- | --- |
| ***Publication record*** |  |
| No. first author papers | (Null)<br>Ethnic group<br>Socioeconomic background<br>Disability, Socioeconomic background<br>Disability<br>Ethnic group, Socioeconomic background |
| No. other author papers | Ethnic group<br>Ethnic group, Sex<br>Disability, Ethnic group |
| ***Job application success*** |  |
| No. postdoc application | Total publications PhD<br>Sex, Total publications PhD<br>Sex, Socioeconomic background, Total publications PhD<br>Age PhD, Sex, Total publications PhD, Age PhD × Sex<br>Disability, Total publications PhD<br>Socioeconomic background, Total publications PhD<br>Disability, Sex, Total publications PhD<br>Sex, Total publications PhD, Sex × Total publications PhD |
| ***Types of contract*** |  |
| Research vs. teaching & research | Disability, Sexual orientation, Permanent or not, Sex,<br>Socioeconomic background, Disability × Sex<br>Disability, Permanent or not, Sex, Socioeconomic background, Disability × Sex<br>Disability, Permanent or not, Total postdocs, Sex,<br>Socioeconomic background, Disability × Sex<br>Permanent or not, Disability × Sex |
|  | Disability, Sexual orientation, Total publications PhD,<br>Permanent or not, Sex, Socioeconomic background, Disability<br>× Sex |
| Permanent or not | Age current, Total postdocs, Total publications PhD<br>Age current, Total postdocs, Sex, Total publications PhD,<br>Total postdocs × Sex<br>Age current, Sexual orientation, Total postdocs, Total<br>publications PhD<br>Age current, Sex, Total postdocs, Total publications PhD<br>Age current, Disability, Total postdocs, Total publications PhD |
| **Grant success** |  |
| No. grant<br>applications | Socioeconomic background, Year PhD<br>Year PhD<br>Sex, Year PhD<br>Sex, Socioeconomic background, Year PhD<br>Sexual orientation, Year PhD<br>Sexual orientation, Socioeconomic background, Year PhD<br>Disability, Socioeconomic background, Year PhD |
| **Reported barriers** |  |
| Reported barrier or<br>not | Sexual orientation, Sex, Socioeconomic background, Year<br>PhD<br>Sexual orientation, Sex, Socioeconomic background<br>Disability, Sexual orientation, Sex, Socioeconomic<br>background, Year PhD |

## References

1. Herring C (2009) Does diversity pay? Race, gender, and the business case for diversity. Am Sociol Rev 74: 208–224.

2. Rohner U, Dougan B (2012) Gender diversity and corporate performance. Credit Suisse Research Institute, Zurich.

3. Herring C (2017) Is diversity still a good thing? Am Sociol Rev 82: 868–877.

4. Hunt V, Prince S, Dixon-Fyle S, Yee, L (2018) Delivering through diversity. McKinsey and Company. Available at https://www.mckinsey.com/~/media/McKinsey/Business%20Functions/Organization/Our%20Insights/Delivering%20through%20diversity/Delivering-through-diversity_full-report.ashx.

5. Valantine HA, Collins FS (2015) National Institutes of Health addresses the science of diversity. Proc Natl Acad Sci USA 112: 12240–12242.

6. Herring C (2014) Diversity and departmental rankings in chemistry. Careers, Entrepreneurship, and diversity: Challenges and opportunities in the global chemistry enterprise, eds HN Cheng, S Shah, ML Wu (American Chemical Society, Washington), pp 225–236.

7. Campbell LG, Mehtani S, Dozier, ME, Rinehart J (2013) Gender-heterogenous working groups produce higher quality science. PLOS One 8: e79147.

8. AlShebli BK, Rahwan T, Woon WL (2018) Ethnic diversity increases scientific impact. Preprint available at https://arxiv.org/abs/1803.02282.

9. Pell AN (1996) Fixing the leaky pipeline: Women scientists in academia. J Anim Sci 74: 2843–2848.

10. Sugimoto CR, Lariviere V, Ni CQ, Gingras Y, Cronin B. 2013. Global gender disparities in science. Nature 504: 211–213.

11. UNESCO Institute for Statistics. 2017. Women in science. Available at http://uis.unesco.org/apps/visualisations/women-in-science/.

12. Käfer J, Betancourt A, Villain AS, Fernandez M, Vignal C, Marais GAB, Tenaillon MI (2018) Progress and prospects in gender visibility at SMBE Annual Meetings. Genome Biol Evol 10: 901–908.

13. Edwards HA, Schroeder J, Dugdale HL (2018) Gender differences in authorships are not associated with publication bias in an evolutionary journal. PLoS ONE 13: e0201725.

14. Holman L, Stuart-Fox D, Hauser CE (2018) The gender gap in science: How long until women are equally represented? PLoS Biol 16: e2004956.

15. Heward C, Taylor P, Vickers R (1997) Gender, race and career success in the academic profession. J Furth High Educ 21: 205–218.

16. Fang D, Moy E, Colburn L, Hurley J (2000) Racial and ethnic disparities in faculty promotion in academic medicine. JAMA 284: 1085–1092.

17. Ng TWH, Eby LT, Sorensen KL, Feldman DC (2005) Predictors of objective and subjective career success: a meta-analysis. Pers Psychol 58: 367–408.

18. Paul B (2016) Diversity, funding, and grassroots organizing. Science: DOI: 10.1126/science.caredit.a1600124.

19. Bhatt W (2013) The little brown woman: Gender discrimination in American medicine. Gender Soc 27, 659–680.

20. Fournier AMV, Bond AL (2015) Volunteer field technicians are bad for wildlife ecology. Wildlife Soc B 39: 819–821.

21. Malcolm SM, Hall PQ, Brown JW (1976) The double bind: The price of being a minority woman in science. American Association for the Advancement of Science, Washington.

22. Ong M, Wright C, Espinosa L, Orfield G (2011) Inside the double bind: A synthesis of empirical research. Harvard Educ Rev 81: 172–209.

23. Armstrong MA, Jovanovic J (2017) The intersectional matrix: Rethinking institutional change for URM women in STEM. J Divers High Educ 10: 216–231.

24. Stout JG, Wright HM (2016) Lesbian, gay, bisexual, transgender, and queer students’ sense of belonging in computing: An intersectional approach. Comput Sci Eng 18: 24–30.

25. National Academy of Sciences, National Academy of Engineering, & Institute of Medicine (2011) Expanding underrepresented minority participation: America’s science and technology talent at the crossroads. The National Academies Press, Washington.

26. R Core Team (2018) R: A language and environment for statistical computing. Vienna, Austria: R Foundation for Statistical Computing. Available at https://www.r-project.org/.

27. Bartoń K (2014) MuMIn: Multi-model inference. [R package version 1.42.1].

28. Hurvich CM, Tsai CL (1989) Regression and time series model selection in small samples. Biometrika 76: 297–307.

29. Burnham KP, Anderson DR (2002) Model selection and multimodel inference: A practical information-theoretic approach. Springer-Verlag, New York.

30. Gelman A (2008) Scaling regression inputs by dividing by two standard deviations. Stat Med 27: 2865–2873.

31. Grueber CE, Nagakawa S, Laws RJ, Jamieson IJ (2011) Multimodel inference in ecology and evolution: challenges and solutions. J Evol Biol 24: 699–711.

32. Cashmore A (2009) Reward and Recognition of Teaching in Higher Education: Institutional Policies and their Implementation (No. 2). The Higher Education Academy and GENIE Centre for Excellence in Teaching and Learning, University of Leicester.

33. Feinerer I, Hornik K (2018) tm: Text Mining Package. R package version 0.7-6. Available at https://CRAN.R-project.org/package=tm.

34. Feinerer I, Hornik K, Meyer D (2008) Text Mining Infrastructure in R. Journal of Statistical Software 25(5): 1–54. Available at http://www.jstatsoft.org/v25/i05/.

35. Lott B (2002) Cognitive and behavioural distancing from the poor. Am Psychol 57: 100–110.

36. Gardner SK, Holley KA (2011) “Those invisible barriers are real”: The progression of first-generation students through doctoral education. Equity Excell Educ 441: 77–92.

37. Palepu A, Carr PL, Friedman RH, Amos H, Ash AS, Moskowitz MA (1998) Minority faculty and academic rank in medicine. JAMA 280: 767–771.

38. Alexander CE, Arday J (2015) Aiming higher: Race, inequality and diversity in the academy. Runnymede Trust, London.

39. Rollock N (2019) Staying power: the career experiences and strategies of UK black female professors. University and College Union, London.

40. Vasquez MJT, Lott B, García-Vázquez E, Grant SK, Iwamasa GY, Molina LE, Ragsdale BL, Vestal-Dowdy E (2006) Personal reflections: Barriers and strategies in increasing diversity in psychology. Am Psychol 61: 157–172.

41. Kameny RR, DeRosier ME, Taylor LC, Sturtz McMillen J, Knowles MM, Pifer K (2014) Barriers to career success for minority researchers in the behavioural sciences. J Career Dev 41: 43–61.

42. Advance HE (2018) Equality in Higher Education: Statistical report 2018.

43. Eby LT, Allen TD, Evans SC, Ng T, DuBois DL (2008) Does mentoring matter? A multidisciplinary meta-analysis comparing mentored and non-mentored individuals. J Vocat Behav 72: 254–267.

44. van der Weijden I, Belder R, van Arensbergen P, van den Besselaar P (2015) How do young tenured professors benefit from a mentor? Effects on management, motivation and performance. High Educ 69: 275–287.

45. Hadani M, Coombes S, Das D, Jalajas (2012) Finding a good job: Academic network centrality and early occupational outcomes in management academia. J Organ Behav 33: 723–739.

46. Parker M, Welch EW (2013) Professional networks, science ability, and gender determinants of three types of leadership in academic science and engineering. Leadership Quart 24: 332–348.

47. Spurk D, Meinecke AL, Kauffeld S, Volmer J (2015.) Gender, professional networks, and subjective career success within early academic science careers. J Pers Psychol 14: 121–130.

48. Intemann K (2009) Why diversity matters: Understanding and applying the diversity component of the National Science Foundation’s broader impacts criterion. Social Epistimol 23: 249–266.

